# IRE1α is critical for kaempferol induced neuroblastoma differentiation

**DOI:** 10.1101/432369

**Authors:** Ahmad Abdullah, Priti Talwar, Palaniyandi Ravanan

## Abstract

Neuroblastoma is an embryonic malignancy arises out of the neural crest cells of the sympathetic nervous system. It is the most common childhood tumor and well known for its spontaneous regression via the process of differentiation. The induction of differentiation using small molecule modulators such as all trans retinoic acid is one of the treatment strategies to treat the residual disease. In this study, we have reported the effect of kaempferol, a phytoestrogen in inducing differentiation of neuroblastoma cells *in vitro*. Treatment of neuroblastoma cells with kaempferol reduced the proliferation and enhanced apoptosis along with the induction of neuritogenesis. Analysis of the expression of neuron specific markers such as β III tubulin, neuron specific enolase and NRDG1 (N-myc down regulated gene 1) revealed the process of differentiation accompanying kaempferol induced apoptosis. Further analysis on understanding the molecular mechanism of action showed that the activity of kaempferol happened through the activation of the endoribonuclease activity of IRE1α (Inositol requiring enzyme 1 alpha), an endoplasmic reticulum (ER) resident transmembrane protein. The *in silico* docking analysis and biochemical assays using recombinant human IRE1α confirms the binding of kaempferol to the ATP binding site of IRE1α and thereby activating ribonuclease activity. Treatment of cells with the small molecule inhibitor STF083010 which specifically targets and inhibits the endoribonuclease activity of IRE1α showed reduced expression of neuron specific markers and curtailed neuritogenesis. The knock down of IRE1α using plasmid based shRNA lentiviral particles also showed diminished changes in the change in morphology of the cells upon kaempferol treatment. Thus our study suggests that kaempferol induces differentiation of neuroblastoma cells via the IRE1α-XBP1 pathway.

## 1. Introduction

Neuroblastoma the most common solid extra cranial malignant tumor of infants originates from the precursor cells derived from neural crest tissues (Duckett and Koop, 1977; Hoehner et al., 1996). The clinical representation of the disease varies drastically from a mass with no symptoms to a primary tumor causing critical illness by disseminating into the neck, thorax, abdomen or pelvis and stands as the most commonly diagnosed tumor in early childhood (Brodeur, 2003). The heterogeneity of neuroblastoma tumors makes intense chemoradiotherapy as a therapeutic strategy. Therefore, for minimizing the effect of chemoradiotherapy, alternative therapeutic strategies targeting crucial proteins involved in the tumor progression is needed. The derivatives of retinoic acids are widely used clinically after a cytotoxic therapy as differentiating agents for controlling minimal residual disease in high risk neuroblastoma patients (Matthay et al., 1999).

Retinoic acid, a derivative of vitamin A has been shown to promote differentiation of embryonic, neural and mesenchymal stem cells (Jones-Villeneuve et al., 1982; Takahashi et al., 1999; Wichterle et al., 2002; Zhang et al., 2006). All trans retinoic acid (ATRA) (Watabe et al., 2002), 9-cis RA and 13-cis RA (Chow et al., 1991) are the well-known RA isomers that have been shown to induce differentiation and used in the treatment of neuroblastoma. Resistance to retinoic acid therapy is commonly observed *in vitro* as well as clinically, due to the mutations in RAR gene or RAR receptor (Pemrick et al., 1994). In addition, trichostatin A, inhibitor of HDAC, kenpaullone and SB-216763 has also been reported to induce neuritogenesis *in vitro* (Balasubramaniyan et al., 2006; Halder et al., 2015; Lange et al., 2011). Recent studies reported alectinib (Anaplastic Lymphoma Kinase inhibitor) and YK-4-279 (mitosis disruptor) as promising candidates for neuroblastoma treatment by emphasizing the lack of effective therapies at present in clinics (Kollareddy et al., 2017; Lu et al., 2017). Inhibitors of HDAC8, Sphingadienes (a growth inhibitory sphingolipid from soy) and p-dodecylaminophenol have been recently reported to have therapeutic efficacy by reducing tumorogenesis *in vitro* and *in vivo* in mice models (Rettig et al., 2015; Takahashi et al., 2017; Zhao et al., 2018b).

IRE1α is an ER resident protein involved in the UPR pathway. It has a luminal N-terminal domain inside ER lumen and a two functional enzymatic cytoplasmic domains having kinase and endoribonuclease activity. The activation IRE1α by the accumulation of misfolded proteins or by other physiological stimuli not limiting to lipid perturbation, viral infection or immune response (Abdullah and Ravanan, 2018b) which leads to the initiation of its kinase or endoribonuclease activity. The kinase activity is known for the recruitment of TRAF2 protein thereby the activation of JNK (c-Jun N-terminal kinase) mediated signalling cascade driving cells towards an apoptotic response is elicited (Urano et al., 2000). The ribonuclease activity of IRE1α can cleave multiple RNA substrates involved in diverse cell response pathways with Xbp1 mRNA being the major substrate (Niwa et al., 2005). Spliced form of Xbp1 mRNA get translated into an active transcription factor XBP1s and involved in the transcription of variety of genes (Acosta-Alvear et al., 2007a). The mouse embryos lacking IRE1α died after 11.5–14.5 days post-coitum emphasizing the role of IRE1α in the developmental stages of the organism (Iwawaki et al., 2009). Moreover, the IRE1α-XBP1 pathway of gene activation is known to be a critical factor involved in the differentiation of plasma cells, osteoblast and adipocytes (Reimold et al., 2001a; Sha et al., 2009a; Tohmonda et al., 2011).

Kaempferol is a phytoestrogen belongs to the class of flavonoids and known to exhibit antioxidant, anti-inflammatory and anti-cancer activity (Wang et al., 2018). In our attempt for identifying an inhibitor against ER stress induced cell death, we found kaempferol to rescue mammalian cells from apoptosis induced by multiple cell death inducers by acting as an allosteric inhibitor for executioner caspases (Abdullah and Ravanan, 2018a). While performing the experiments we observed the change in morphology of IMR32 cells, a human neuroblastoma cell line with neurite like outgrowths upon prolong incubation in the presence of kaempferol.

In this study, we have investigated the effect of kaempferol on IMR32 and Neuro2a neuroblastoma cell lines. Observations made in our experiments suggest that kaempferol mediated differentiation of neuroblastoma cells happens via the activation of IRE1α endoribonuclease activity. Since Kaempferol is a dietary polyphenol and cohort studies had been performed for understanding its effect on various other cancers (Cui et al., 2008; Gates et al., 2009; McCann et al., 2005), it could also be explored for its effect on neuroblastoma in clinical trials.

### 2. Materials and Methods

### 2.1 Cell culture and reagents

IMR32, human neuroblastoma cell line and Neuro2A, mouse neuroblastoma cells were obtained from NCCS, Pune, India. Cells were maintained in DMEM containing 10% FBS (Himedia), 1X glutamine and 1X antibiotic and antimycotic solution at 37 °C and 5%CO_2_. Kaempferol (HPLC purity≥98.8%) was obtained from Natural remedies (Bangalore, India). Diarylpropionitrile (DPN) and 4-[2-Phenyl-5,7-*bis*(trifluoromethyl)pyrazolo[1,5-*a*] pyrimidin-3-yl] phenol (PHTPP) were obtained from Alfa Aesar (Hyderabad, India). APY29 was purchased from MedChem express, USA. STF083010 was purchased from Cayman chemicals, USA.

### 2.2 Cell viability assay

MTT assay was performed for analyzing the cytotoxic effect of kaempferol on IMR32 and Neuro2A cell lines. Briefly, 0.5 × 10^4^ cells were seeded per well in a 96 well plate and treated with kaempferol for 48 h, 72 h and 96 h. After the desired incubation time, 20 µl of 5 mg/ml MTT (Himedia) was added to each of the well and incubated for 3 hours. The terazolinium salt formed by the live cells was dissolved by adding 200 µl of DMSO, after the removal of media. The purple color formed was read at 570 nM using ELISA plate reader.

### 2.3 Cell proliferation assay

For trypan blue dye exclusion assay, 0.5 × 10^5^ cells per well was seeded in a 24 well plate. Cells were treated with kaempferol (25, 50 and 100 µM), ATRA (25 µM) and DMSO (vehicle) for a period of 48 h, 72 h and 96 h. Cell counting was done manually using hemocytometer, by staining the cells with trypan blue solution prepared in 0.4% methanol. Live cells that did not take up the dye were counted to quantitate the rate of proliferation under control and treated conditions.

### 2.4 Differentiation experiments

For differentiation experiments, neuroblastoma cell lines were seeded at a density of 3×10^5^ cells in 60mm dish and treated with 50 µM of kaempferol. Induction was given once for a continuous time of 96 hours without any replacement of media. ATRA at a concentration of 25 µM was used as a positive control to compare the morphological changes; images were taken using phase contrast microscope at specified time points.

### 2.5 Stable lentiviral transductions

The IRE1α (sc-40705-V) shRNA lentiviral particles was purchased from Santa Cruz Biotechnology (Santa Cruz, CA, USA) contain expression constructs encoding target-specific shRNA designed to specifically knock down human IRE1α gene expression. Scrambled shRNA contain lentiviral particles (sc-108080; Santa Cruz Biotechnology) that will not lead to specific degradation of any cellular mRNA was used as negative control. Lentiviral transduction was performed in the presence of polybrene (5 µg/ml, Sigma-Aldrich), and stable transductants were generated via selection with puromycin (1 µg/ml, Sigma-Aldrich) according to the manufacturer’s instructions.

### 2.6 Immunocytochemistry and microscopy

Immunocytochemistry was carried out as described in the Cell signaling Technology (CST) guide. Briefly, 2.5 × 10^4^ cells/well were seeded in a 24 well plate. After overnight incubation, cells were treated with the compounds for indicated time. The cells were then washed with 1XPBS and fixed with 4% Formaldehyde for 15 min. The cells were washed again with 1XPBS and incubated with 0.5% skimmed milk containing 0.3% tritonX100 for blocking nonspecific binding, for 1 hour. Immunostaining was done using β-III tubulin (CST #5568) and Synaptophysin (Novus #NB2-25170SS) specific primary antibodies at 4°C with overnight incubation followed by 1 hour incubation with the secondary antibody, anti-rabbit Alexa Flour 594 (#8889, Cell Signaling Technologies, USA). Phase contrast and fluorescence images of cells under treated and non-treated conditions were captured using EVOS FLoid imaging station (Thermo Fisher, USA) with 20X fluorite objective and LED light cubes containing hard coated filters (blue and red). For a particular protein expression study, imaging parameters were kept constant throughout the imaging and no image modifications were done post imaging.

### 2.7 Gene expression analysis

Total RNA isolation was performed using RNA isoplus (Takara, India). RNA quantification was performed and 2 µg of total RNA was used for cDNA conversion using Primescript RT reagent kit (Takara, India). Quantitative-PCR (qRT-PCR) was performed using SYBR green reagent (Takara, India) using Applied Biosystems Quant studio 3 real time PCR machine. Human 18s rRNA was used as an internal control for normalizing the gene expression. The fold induction of mRNA levels was calculated by 2^−ΔΔ^*^CT^method*. The list of primers used in this study is tabulated (Table 1).

**Table 1:**
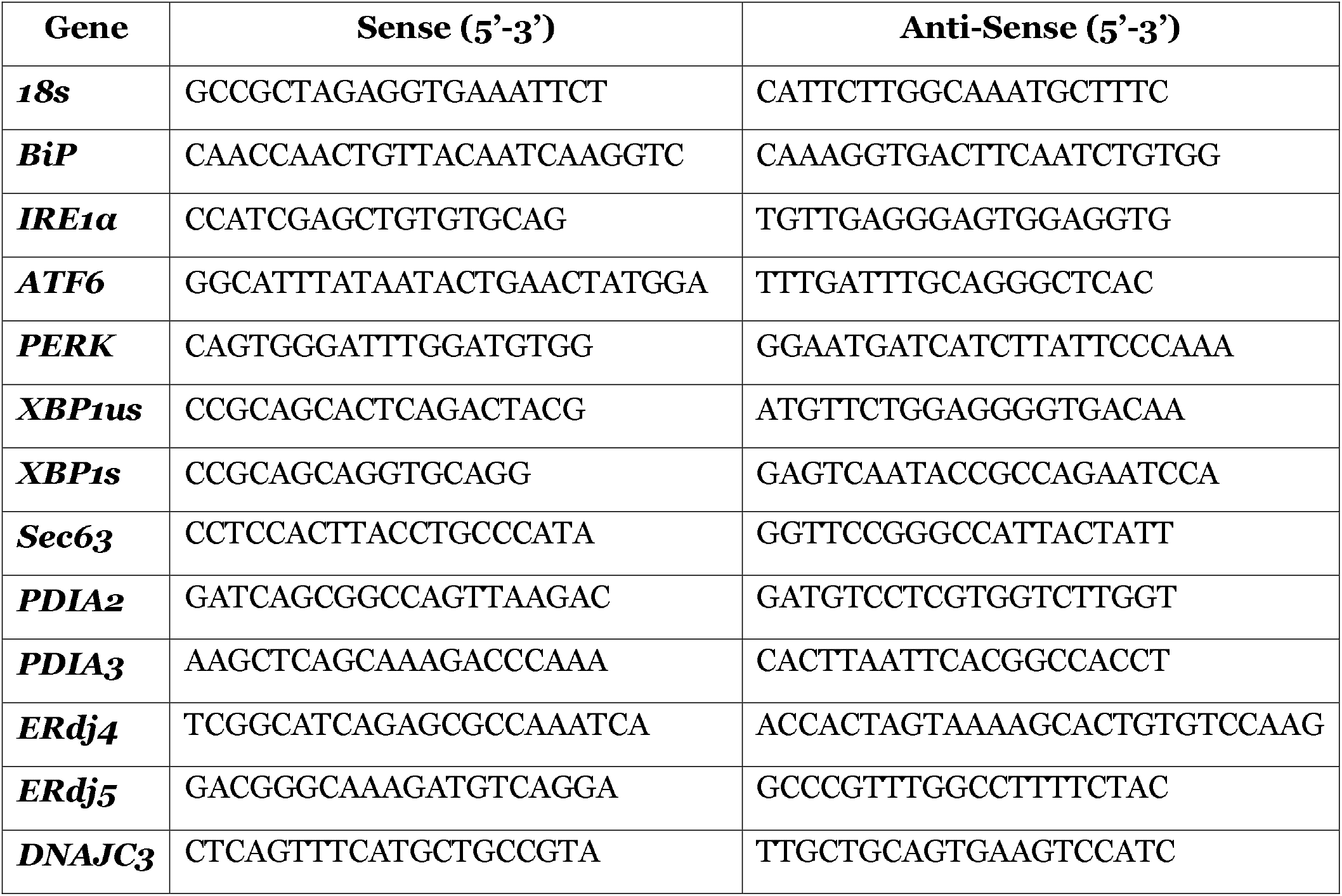
List of primers used in this study.

### 2.8 Immunoblotting

Immunoblotting was carried out as described earlier by Ravanan et al (Ravanan et al., 2011). In brief, cells were plated in 60 mm dishes at the density of 1.5 × 10^5^ cells/dish. After overnight incubation, cells were treated with mentioned compounds. At 24, 48, 72 and 96 h time points after the treatment, cells were washed with ice cold PBS and lysed with RIPA buffer containing 1X Protease inhibitor cocktail (Sigma Aldrich). Total protein concentrations were quantified by Folin’s assay. Immunoblotting analysis was done by loading 50 µg of protein in each lane for SDS-PAGE. Antibodies used in this study were obtained from Cell Signalling Technology, USA (Beta actin#8457, N-myc#9405, IRE1α#3294 and XBP1s#12782), Novus biological, USA (NDRG1#NBP1-32074, ERα#NB120-3577 and ERβ#NB100-92166) and ABclonal, USA (NSE #A3118 and GRP78 #A0241). Anti-rabbit, HRP linked secondary antibody from Cell Signalling Technology (#7074) was used for chemiluminescence based detection.

### 2.9 Methylene blue staining and neurite counting

Cells were seeded in 35 mm dish at a density of 1.5 × 10^5^ cells and differentiation was induced with kaempferol (50 µM) or APY29 (250 nM). Wherever STF083010 (50 µM) is used as inhibitor, cells were pre-treated with STF083010 for 90 min and then treated with the mentioned test compounds. After 96 h of incubation, cells were fixed using 4% formaldehyde for 15 min and stained using 0.2% methylene blue solution in methanol for 30 min. After repeated washes for unbound stain removal, images were taken using monochrome phase contrast camera of EVOS FLoid imaging station (Thermo Fischer). Total number of cells and cells bearing 2 or more neurites were counted for 10 random focuses and represented as percentage of cells in the graph (Chaudhari et al., 2017b).

### 2.10 Kinase inhibition studies

Recombinant human IRE1α enzyme was obtained from SignalChem (#E31-11G). ADP Glo kinase assay (Promega) was performed as per the manufacturer’s protocol to confirm the inhibitory mechanism of kaempferol. Briefly, 50 ng of enzyme was used with 10 µM or 100 µM of ATP with different concentrations of kaempferol in a white flat bottom 96 well plate. Staurosporine (Sigma Aldrich) a pan kinase inhibitor was used as positive control. The luminescence produced by the conversion of dephosphorylated ADP to ATP was measured using the luminometer (Berthold).

### 2.11 *In silico* docking analysis

Studies to understand the mode of binding of kaempferol to IRE1α was carried out using AutoDock suite 4.0. Crystal structure of the human IRE1α bound with ADP (PDB ID: 3P23) was used as the target protein. Ligands were obtained from Pubchem as .sdf format and converted to .pdb using smiles translator. Polar hydrogen atoms were assigned for the macromolecule prior docking. Gasteiger charges for ligands and Kollman charges for the receptor molecule was added using AutoDock tools 1.5.6. Throughout the study, the macromolecule was kept rigid with rotatable bonds assigned for the ligands. The grid center made on the macromolecule covering the entire surface of the protein acted as search space. AutoGrid 4.0 was used to produce the map files of the flexible atoms and docking parameter file (DPF) was generated using Lamarckian Genetic algorithm of AutoDock 4.0. Validation of the docking study was carried out by re-docking the ADP with the solved crystal structure of kinase domain of IRE1α. The re-docked complex was compared to the known crystal structure of the ligand-macromolecule complex. Binding energies of the best docked pose of the docked complexes were calculated based on torsional energy, H-bonding, non-bonded interactions and desolvation energies. LigPlot^+^ v. 1.4 was used to study the interactions at 2D level while UCSF Chimera 1.10.2 and PyMol V. 1.7.4 were used in visualizing 3D interactions.

### 2.12 Statistical analysis

All data are presented as the Standard error of mean (S.E.M) of at least three independent experiments. Statistical comparisons were performed using one-way/two-way analysis of variance (ANOVA) followed by Bonferroni’s multiple comparison test (GraphPad Prism, version 5.0) and p-value of less than 0.05 was considered statistically significant.

## 3. Results

### 3.1 Kaempferol induces neuroblastoma differentiation and decreases cell viability

To understand the effect of kaempferol on neuroblastoma cells, we employed IMR32 and Neuro2a cell lines, well-established in vitro models that have been used widely for understanding neuroblastoma differentiation. IMR32 and Neuro2a cells were incubated in the presence of kaempferol (50 µM) for 4 days, with the treatment in the presence of ATRA (25 µM) served as a positive control. After 48 h of incubation, we observed morphological changes including neurite outgrowth associated with decreased proliferation and viability. To confirm whether the neurite outgrowth is due to cellular differentiation, we performed immunocytochemistry to analyze the expression of β-III tubulin, a neuronal differentiation marker. The significant increase in the expression level of β-III tubulin in the kaempferol and ATRA treated cells compared to the controls confirmed the neurite outgrowth to be due to the cellular differentiation (Fig. 1A). A further assessment on the viability of the cells using MTT assay and trypan blue dye exclusion assay revealed that kaempferol decreased the viability of both IMR32 and Neuro2a treated cells (Fig. 1B and Supplement 1A and B). An additional quantitation of the number of cells bearing 2 or more neurites indicated that an incubation period of 4 days is enough for kaempferol to effectively induce a significant change in the morphology (Fig. 1C and D), while in ATRA treated cells there was lesser morphological changes in terms of neurite extensions when compared to kaempferol treated cells. ATRA had been shown to require 10–14 days for a significant change in morphology (Pence and Shorter, 1990), while kaempferol induced it in 4 days of treatment. Complementarily, we observed an increase in the mRNA and protein levels of certain neuron specific markers genes like synaptophysin (*SYP*), neuron specific enolase (*NSE*) and N-myc down regulated gene 1 (*NDRG1*) (Fig. 2A and B). Although the mRNA transcripts of *NSE* and *NDRG1* in ATRA treated condition didn’t show significant up-regulation, the protein levels were up-regulated during day3 and day4 of differentiation. The expression of N-myc, at mRNA level, was down-regulated in kaempferol treated conditions while it was significantly increased in ATRA treated conditions (Fig. 2A). In contrast, the down-regulation of *N-myc* observed at the mRNA levels was not exactly the same at the protein level as we observed no such drastic reduction in N-myc protein levels (Fig. 2B). The reason for this could be the accumulation of N-myc protein due to protein stabilization rather than transcription of the gene (Choi et al., 2010). Moreover, the increase in the expression of N-myc protein is shown to be a required criterion for the onset of differentiation (Guglielmi et al., 2014). However, we observed an increase in NDRG1 expression levels with a decrease in the expression level of N-myc protein between day 2 and day 4 (Fig. 2B); a phenomenon reported to occur during neuroblastoma differentiation (Kang et al., 2006). Altogether, our data suggest that kaempferol promotes differentiation and cell death in neuroblastoma cells.

**Figure 1:**
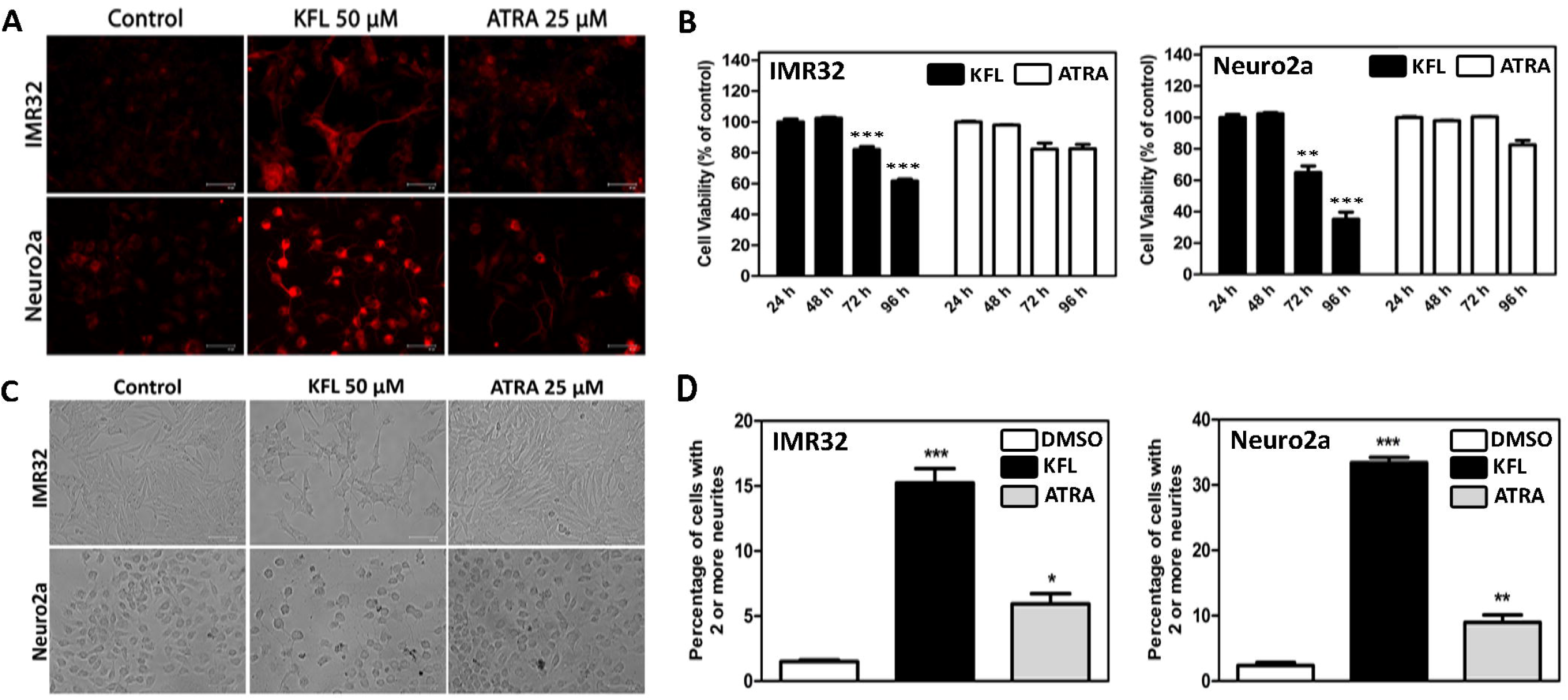
Kaempferol induces differentiation of neuroblastoma cells. **A.** Immunocytochemistry images showing the expression of β III tubulin expression in IMR32 and Neuro2a cells treated with kaempferol (KFL) and ATRA for a time period of 96 h. Representative images of experiments performed more than three times. Scale bars = 67 µm. **B.** Cell death analysis using MTT cell viability assay with Kaempferol (KFL) 50 µM and ATRA 25 µM. The average ±SEM from 3 independent experiments performed in triplicates. (**p < 0.01; ***p < 0.001; two-way ANOVA) **C.** Phase contrast images showing the change in morphology and neurite outgrowth with Kaempferol (KFL) and ATRA treatment in IMR32 and Neuro2a cells for 96 h. Representative images of experiments performed more than three times. Scale bars = 67 µm. **D.** Graphs representing the number of cells bearing 2 or more neurites in IMR32 and Neuro2a cells after 96 h of incubation with Kaempferol (KFL) or ATRA. The average ±SEM from 10 independent counting is shown. (*p < 0.05; **p < 0.01; ***p < 0.001; one-way ANOVA)

**Figure 2:**
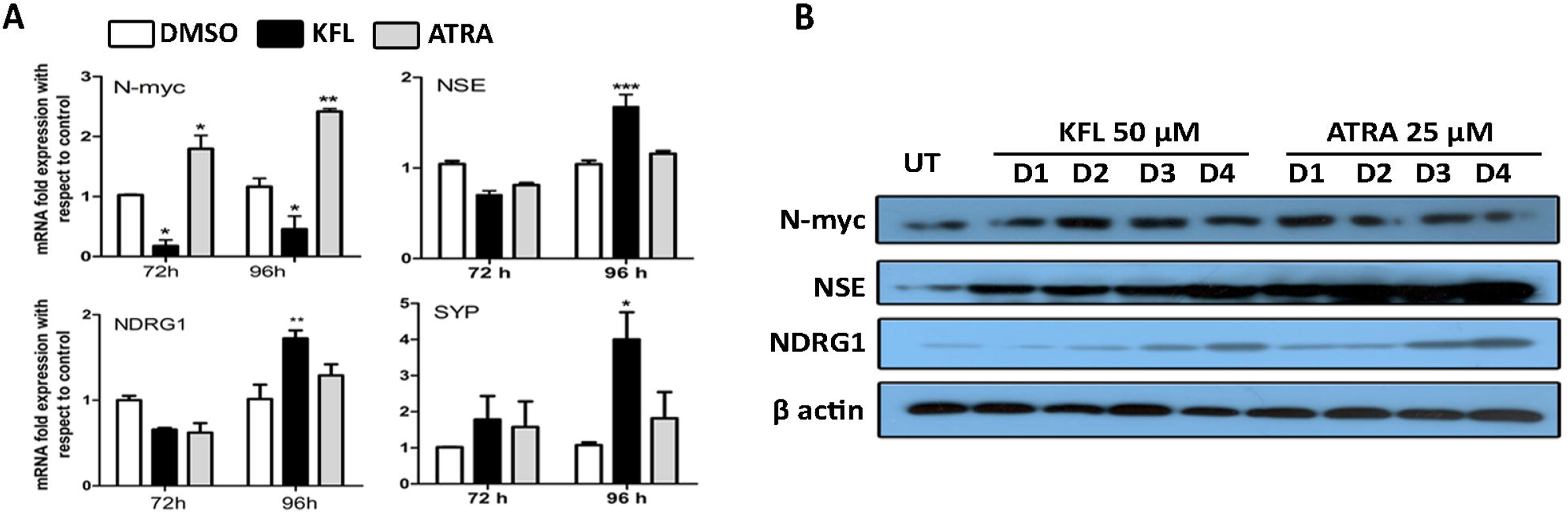
Kaempferol enhances the expression of differentiation markers in IMR32 cells. **A.** Graphs representing the mRNA expression of synaptophysin (SYP), Neuron specific enolase (NSE), NDRG1 and N-myc genes upon Kaempferol (KFL) and ATRA treatment for 72 and 96 h. The average ±SEM from three independent experiments. (*p < 0.05; **p < 0.01; ***p < 0.001; two-way ANOVA) **B.** Western blots performed for NDRG1, NSE and N-myc protein in IMR32 cells showing the expression upon kaempferol (KFL) and ATRA treatment for 4 days (Day1 – Day4).

### 3.2 Kaempferol induced differentiation is independent of the estrogen receptor signaling

It has been previously reported that kaempferol as a phytoestrogen (estrogen receptor agonist) suppresses the proliferation of breast cancer cells and promotes apoptosis (Kim and Choi, 2013). The use of antagonist against estrogen receptors had shown to reverse the biological activity of kaempferol (Wang et al., 2009). Since, ERβ signaling has been shown to be essential for kaempferol mediated biological responses (Kuiper et al., 1998) we examined the expression of ERα and ERβ in IMR32 cells. In our study, we observed that kaempferol substantially instigates the expression of estrogen receptor β (ERβ) compared to the estrogen receptor α (ERα) both at mRNA and protein level in IMR32 cells (Fig. 3A and B). Later on, intending to investigate whether ERβ signaling is involved in the modulation of kaempferol induced neuroblastoma differentiation, we used PHTPP (4-[2-Phenyl-5,7-*bis*(trifluoromethyl)pyrazolo[1,5-*a*]pyrimidin-3-yl]phenol), a selective ERβ antagonist to find out if PHTPP pretreatment interferes with the differentiation process. Interestingly, pretreatment of kaempferol treated cells with PHTPP had no effect both on the formation of neurites and β-III tubulin expression suggesting that kaempferol induced differentiation is independent of ERβ signaling (Fig. 3C). Since there was a basal expression of ERα was observed in IMR32 cells upon kaempferol pretreatment in our study and Wang et al., 2009, have used tamoxifen to reverse the activity of kaempferol (Wang et al., 2009). Therefore we performed an experiment with pretreatment of IMR32 cells with tamoxifen at different concentrations. The pre-treatment of IMR32 cells with tamoxifen an ERα antagonist also did not show any observable effects on the process of differentiation induced by kaempferol (data not shown). In addition, Diarylpropionitrile (DPN), another ERβ agonist, failed to induce differentiation of IMR32 cells. Moreover, pretreatment of PHTPP did not reverse the cell death induced by kaempferol as well (Fig. 3D). Collectively, these findings suggest that kaempferol induced neuroblastoma differentiation and cell death is not mediated by ERβ signaling.

**Figure 3:**
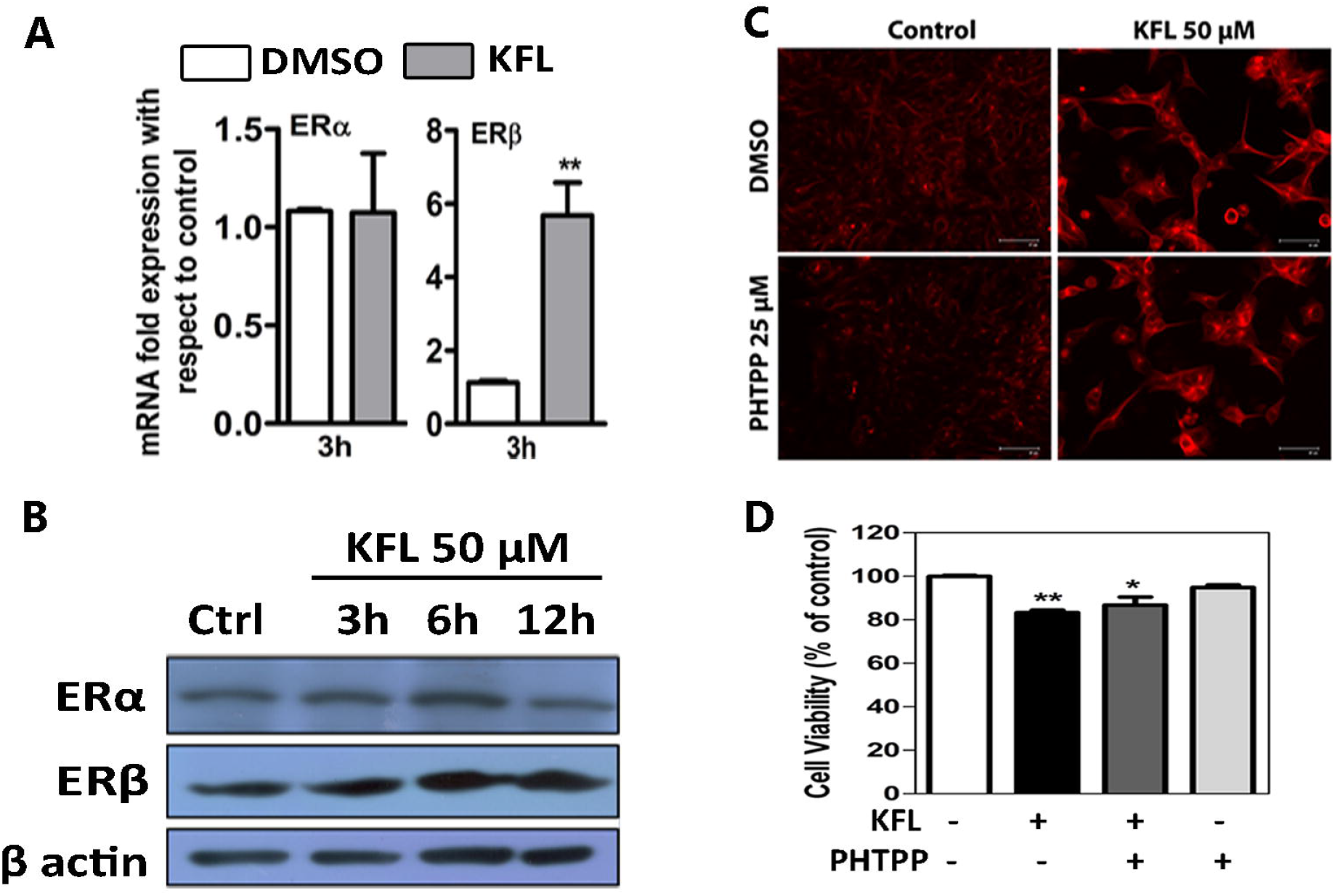
Estrogen pathway independent activity of kaempferol. **A.** mRNA expression levels of estrogen receptor (ER) α and β upon kaempferol (KFL) treatment in IMR32 cells. The average ±SEM from three independent experiments. (*p < 0.05; **p < 0.01; one-way ANOVA) **B.** Western blot analysis showing the regulation of ER α and β expression upon kaempferol (KFL) treatment in IMR32 cells. **C.** Immunocytochemistry experiment showing expression of β III tubulin with kaempferol and with or without PHTPP pretreatment in IMR32 cells. Representative images of experiments performed more than three times. Scale bars = 67 µm. **D.** Cell viability analysis for IMR32 cells using MTT assay showing no effect on the percent viability of cells pretreated with PHTPP (25 µM) in presence of kaempferol (KFL). The average ±SEM from 3 independent experiments performed in triplicates. (*p < 0.05; **p < 0.01; one-way ANOVA)

### 3.3 Kaempferol modulates the expression of IRE1α

Endoplasmic reticulum (ER) stress is known to be associated with various cellular differentiation processes (Ma et al., 2010; Matsuzaki et al., 2015; Wielenga et al., 2015) and kaempferol has been known to induce ER stress in certain cancer cell lines (Guo et al., 2017; Guo et al., 2016; Huang et al., 2010). These research findings prompted us to check the expression level of ER stress marker genes such as *BiP, PERK, IRE1*α *and ATF6* in kaempferol treated IMR32 cells. The results presented in Fig. 4A show that kaempferol significantly increased the expression of *IRE1α,* while that of the other ER stress marker genes was not significantly changed with time (Fig. 4A). IRE1α is an ER transmembrane sensor of unfolded protein response pathway (UPR) with both kinase and endoribonuclease activity (Bork and Sander, 1993). During ER stress, IRE1α activation initiates the splicing of XBP-1 (X-Box binding Protein-1) mRNA which in turn translated into XBP1s protein and involved in transcription of diverse set of genes including genes involved in the UPR pathway (Acosta-Alvear et al., 2007a). Previous studies have reported the involvement of IRE1α-XBP1 pathway in the differentiation of plasma cells (Todd et al., 2009), osteoclasts (Tohmonda et al., 2013) and adipocytes (Sha et al., 2009b) in addition to the unfolded protein response pathway. This led us to hypothesize whether kaempferol might induce the neuroblastoma differentiation by modulating the IRE1α-XBP1 pathway. We then performed protein analysis of IRE1α for kaempferol treated cells and compared it with brefeldin A (a known inducer of ER stress) treated cells. Our results indicated that kaempferol increased the expression of IRE1α but not BiP at protein level. BiP is a molecular chaperone whose increased level is considered as a hallmark of the ER stress process. In contrast, brefeldin A treatment up regulated the levels of IRE1α as well as BiP protein level indicating the onset of strong ER stress response (Fig. 4B). We have previously showed the bioactivity of kaempferol in ameliorating ER stress induced cell death with a reduction in the level of ER stress markers expression in kaempferol pretreated conditions in IMR32 cells (Abdullah and Ravanan, 2018a). Our present study showed that kaempferol increases the expression of IRE1α without activating ER stress in IMR32 cells.

**Figure 4:**
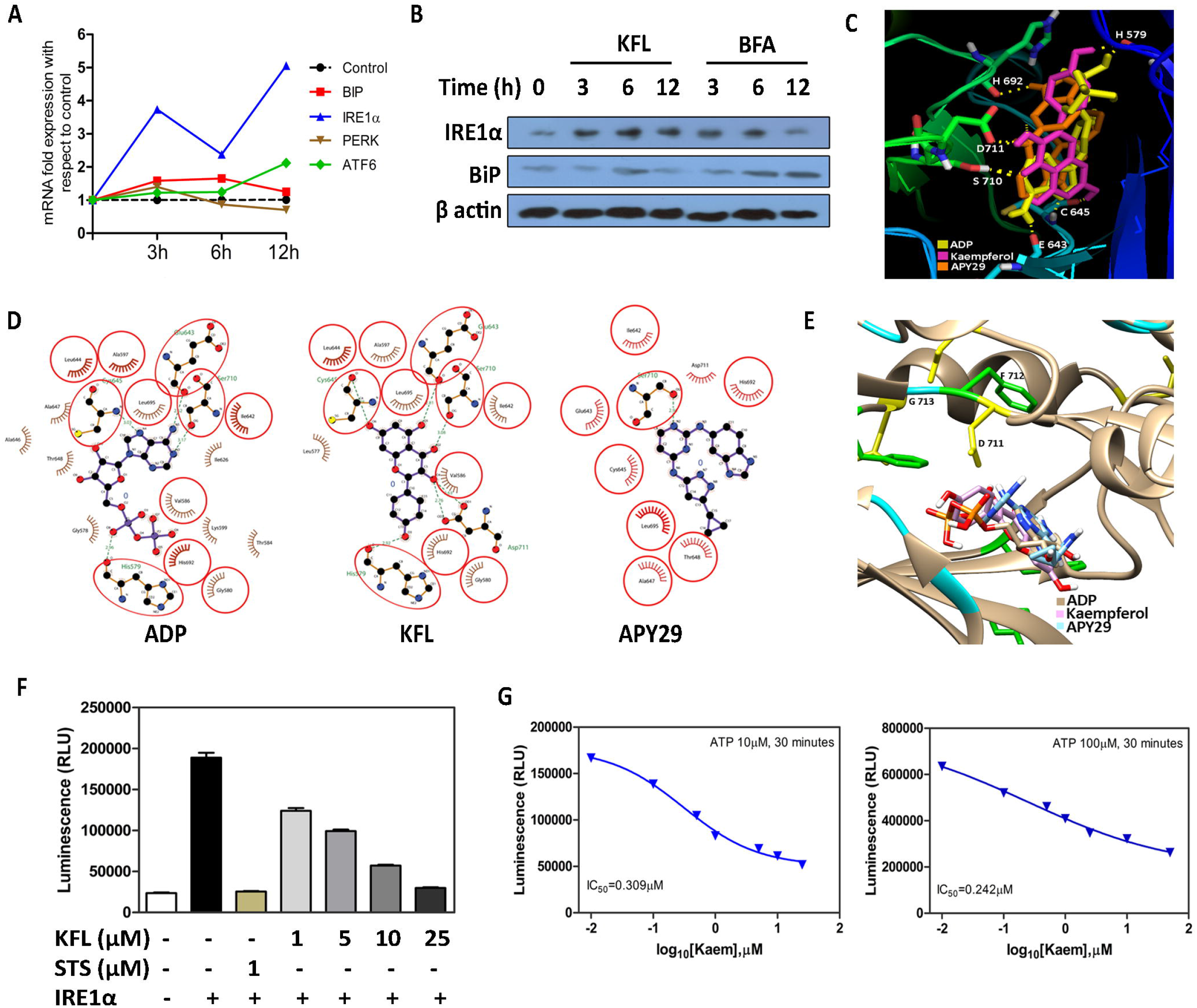
IRE1α modulation by kaempferol via binding to kinase domain. **A.** Gene expression analysis for ER stress marker genes with kaempferol pretreatment in IMR32 cells. The average ±SEM from three independent experiments. (*p < 0.05; ***p < 0.0o1; two-way ANOVA) **B.** Western blot analysis showing the expression of IRE1α and BiP expression on treatment with kaempferol 50 µM (KFL) and Brefeldin A 1 µg/ml (BFA) at 3, and 12 hour time points. **C.** Docked poses of ADP, kaempferol, and APY29 at the nucleotide binding site of human IRE1α kinase domain obtained using Autodock 4.0 and superimposed representation was made using PYMOL. **D.** LigPlot^+^ analysis (2D representation) of the docked complexes of ADP, kaempferol and APY29. Red circles represent the common amino acid residues making interactions with all three compounds. Dashed green lines represent the hydrogen bonding between the amino acid residue and the compound. **E.** Docked poses of ADP, kaempferol and APY29 at DFG-in confirmation (D711, F712 and G713) of human IRE1α (PDB ID: 3P23). Superimposed docked poses and interaction analysis was performed using UCSF chimera. **F.** ADP Glo assay for kinase inhibition by kaempferol (KFL). Staurosporine (STS) was used as a positive control for kinase inhibition activity. Graphs representing the ±SEM of experiment performed in triplicate. **G.** Kinase inhibition assay performed using 10 µM and 100 µM of ATP with same range of concentrations of kaempferol. No significant change in the IC_50_ value shows non-competitive mode of binding of kaempferol to the kinase domain of human IRE1α.

### 3.4 Activation of IRE1α endoribonuclease activity by kaempferol

Quercetin, a flavonoid, has been reported to bind to the Q-site in human-yeast chimeric IRE1 and activates endoribonuclease activity to enhance the cleavage of XBP1 mRNA (Wiseman et al., 2010). Kaempferol also belongs to the flavonoid class of phytocompounds. Considering this, we employed computational docking studies using Autodock to check if kaempferol binds to human IRE1α molecule. Blind docking was performed by selecting the complete surface of the protein molecule. The result showed the kaempferol’s binding affinity towards the kinase pocket of IRE1α (Fig. 4C). The binding of kaempferol was compared to bound states of indigenous activator of IRE1α, ADP and APY29 (known IRE1 kinase inhibitor and endoribonuclease activator) (Fig. 4D). The bound state of ADP in the IRE1α nucleotide binding site is known to activate the endoribonuclease activity of the molecule (Lee et al., 2008). The docked poses analyzed for amino acid residues occupying the bound state of kaempferol showed that the binding of kaempferol to the kinase pocket of the molecule share the similar amino acid residues for interactions as of ADP and APY29 (Fig. 4D). The interactions of kaempferol with Glu 643, Cys 645 and His 692 residues of human IRE1α had been shown for kinase activity inhibition by staurosporine, a pan kinase inhibitor (Concha et al., 2015). APY29 has been known to interact with amino acid residues of kinase domain to activate the endoribonuclease activity of IRE1α (Korennykh et al., 2009). Moreover, the interaction with Cys 645 at the kinase cleft has been reported for APY29 to activate the IRE1α endoribonuclease activity (Wang et al., 2012). The conformation of DFG (Aspartic acid-Phenylalanine-Glycine) motif of kinase domain in IRE1α with an activator for endoribonuclease activity should be in DFG-in conformation rather than DFG-out (Wang et al., 2012). The ADP bound crystal structure of IRE1α (PDB ID: 3P23), with DFG-in motif showing favorable binding affinity for docking with kaempferol suggests its ability to modulate the IRE1α endoribonuclease activity. Moreover, docked pose of kaempferol falls into the same pocket where ADP and APY29 are known to interact with the kinase domain of IRE1α (Fig. 4E). Collectively, the docking analysis showed that kaempferol binds to the kinase domain of IRE1α and could activate the endoribonuclease activity.

To confirm the docking results further, ADP-Glo kinase assay (Promega) was performed using recombinant kinase active human IRE1α. Incubation of kaempferol with IRE1α showed a dose dependent decrease in the kinase activity of IRE1α (Fig. 4F), confirming its binding to the kinase domain of the protein and thereby inhibiting IRE1α’s kinase activity. The kinase inhibition assay performed by increasing the concentration of the substrate ATP from 10 µM to 100 µM, suggested a non-competitive mode of inhibition favored by kaempferol (Fig. 4G), which again correlates with the previously reported weak binding of quercetin (Wiseman et al., 2010) to the nucleotide binding site of human-yeast chimeric IRE1 protein.

### 3.5 IRE1α knockdown reduces neuroblastoma differentiation

To further confirm the role of IRE1α-XBP1 pathway in neuroblastoma differentiation, we established stable IRE1α knock down clones of IMR32 cells using lentiviral particles containing IRE1α shRNA (4 target specific constructs that encodes 19–25 nt shRNA). Successful knock-down of IRE1α at mRNA and protein level was confirmed by qRT-PCR and immunoblotting respectively (Fig. 5A and B). The functional involvement of IRE1α in neuroblastoma differentiation is confirmed, when neither of kaempferol and APY29 exposure triggered β-III tubulin expression in IRE1α knockdown IMR32 cells (Fig. 5C). Additionally, to compare the expression of neuronal differentiation markers (NDRG1 and NSE) between IRE1α knockdown cells and control shRNA transduced cells, we performed immunoblotting analysis. We observed a decrease in the expression level of these markers in IRE1α knockdown cells when compared to the control shRNA transduced cells (Fig. 5D). Also we noticed a reduction in the XBP1s protein level in IRE1α shRNA treated conditions on day4, with a reduced level of differentiation observed. To confirm the activity of endoribonuclease activity in inducing the differentiation of neuroblastoma cells, we inactivated the endoribonuclease activity of IRE1α by using STF083010, a small molecule inhibitor. STF083010 is a specific inhibitor which binds to the endoribonuclease domain of IRE1 and inhibits the cleavage of XBP1 mRNA, while the kinase activity is left intact (Papandreou et al., 2011). The extent of differentiation repression in IRE1α knockdown cells, in the presence of kaempferol or APY29 was further exemplified by the addition of STF083010 (the IRE1 endoribonuclease inhibitor), to these cells (Fig. 5E). Similarly, the number of cells with 2 or more neurites was significantly less in IRE1α knockdown condition upon kaempferol or APY29 treatment, which furthermore decreased upon pretreatment with STF083010 (Fig. 5F). These results validate the role of IRE1α’s endoribonuclease activity in kaempferol induced neuroblastoma differentiation.

**Figure 5:**
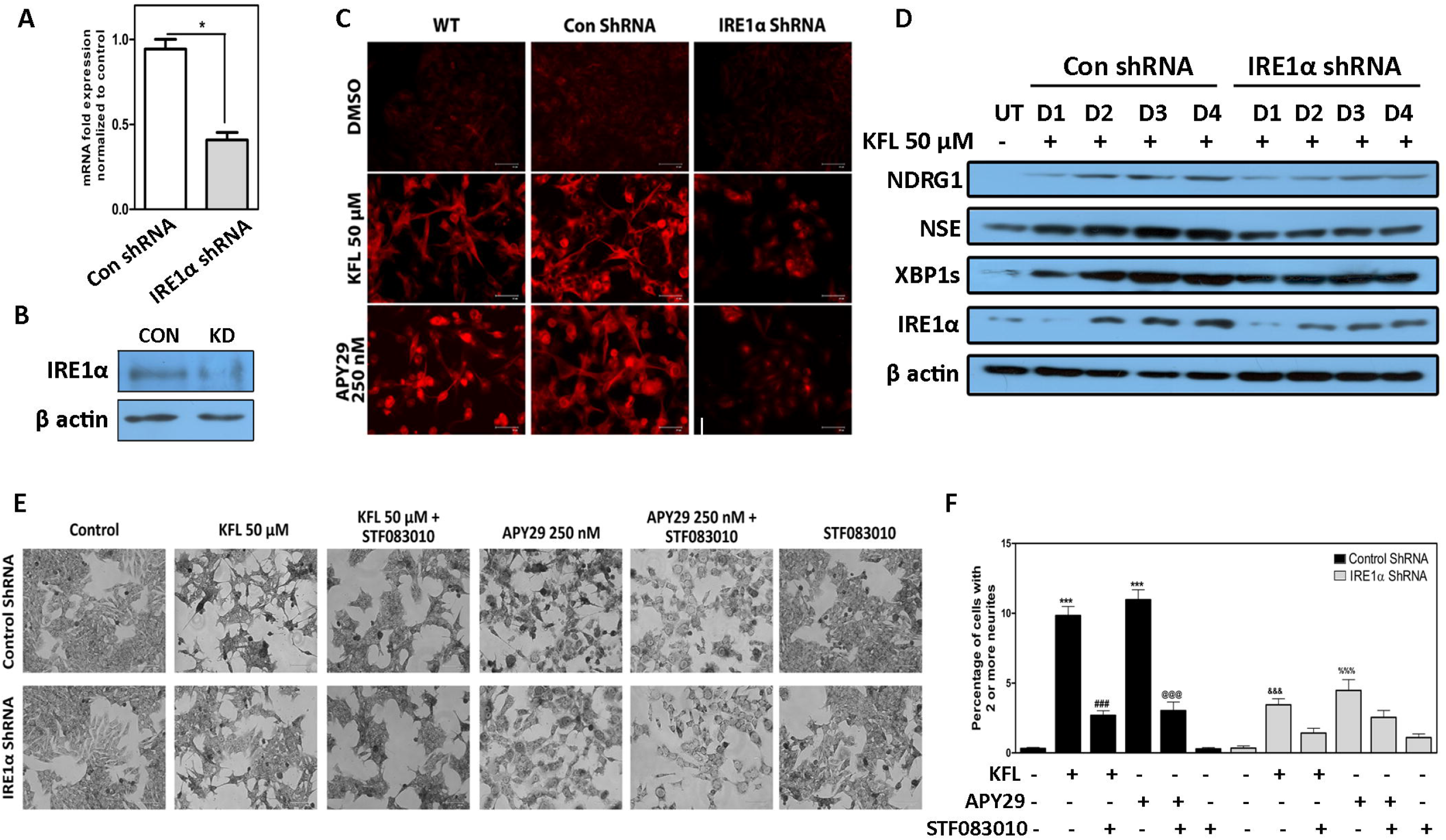
IRE1α knockdown reduces differentiation of IMR32 neuroblastoma cells. **A.** Confirmation of knock down of IRE1α in IMR32 cells at mRNA level. The graph represents average ±SEM of experiment performed in triplicates. (*p < 0.05; one-way ANOVA) **B.** Western blot analysis showing IRE1α knockdown in IMR32 cells. IRE1α shRNA treated cells (KD) showing reduced expression than control shRNA (CON) treated cells. **C.** Immunocytochemistry for expression of β III tubulin showing reduced expression in IRE1α knockdown cells in the presence of kaempferol and APY29. Representative images of experiments performed more than three times. Scale bar = 67 µm. **D.** Western blot analysis for the expression of neuronal markers showing reduced expression in IRE1α knockdown cells. Reduction in the expression of XBP1s levels was also observed in IRE1α knockdown conditions. **E.** Phase contrast methylene blue stained images of IMR32 cells showing change in morphology and neurite outgrowth upon treatment with kaempferol (KFL) and APY29 for 96 h in control shRNA and IRE1α shRNA treated cells. Pretreatment with STF083010 (50 µM) inhibits the process of neuritogenesis further in IRE1α shRNA treated cells. Representative images of experiments performed more than three times. Scale bar = 100 µm. **F.** Graphs representing the number of cells with 2 or more neurite per cell in control shRNA and IRE1α shRNA treated IMR32 cells after 96 h incubation with kaempferol and APY29. STF083010 pretreatment reduces the number of neurite bearing cells in both kaempferol (KFL) and APY29 treated conditions further. The average ±SEM from 10 independent counting is shown.* represents comparison between control and kaempferol/APY29 treated cells; # and @ represents comparison between kaempferol/APY29 treated cells with STF083010 pretreated cells; $ and % represents comparison between kaempferol and APY29 treated conditions in control shRNA and IRE1α shRNA treated cells respectively (*p < 0.05; **p < 0.01; ***p < 0.001; two-way ANOVA).

### 3.6 XBP1s drives differentiation of neuroblastoma cells

Activated IRE1α splices the *XBP1* mRNA which then gets translated into an active transcription factor that regulates diverse set of genes involved in the regulation of various cellular processes including genes related to ER stress, differentiation, cell cycle arrest and immune response (Acosta-Alvear et al., 2007b). The ratio of spliced to unspliced form of XBP1 mRNA corresponds to the extent of IRE1α activation. Small molecule modulators binding to the nucleotide binding site of IRE1α can induce the endoribonuclease activity in the absence of ER stress (Han et al., 2009).

Therefore, we inclined to check the expression level of spliced *XBP1* mRNA by qRT-PCR. IMR32 cells were treated with kaempferol or brefeldin A. Comparing the mRNA expression of *XBP1* unspliced to spliced form, we observed an enhanced presence of XBP1 spliced mRNA levels upon kaempferol treatment. The increase in the expression of *XBP1* (unspliced) mRNA correlates with the enhanced expression of ATF6 (Tsuru et al., 2016). Whereas, this was not the case for brefeldin A treated cells as they displayed a decreased expression level of XBP1 unspliced form (Fig. 6A). To confirm that XBP1s are transcriptionally active, we analyzed its downstream targets such as *SEC63, PDIA2, PDIA3, DNAJC3, ERDJ4 and ERDJ5* mRNA level by qRT-PCR, whose regulation by the transcription factor XBP1s was validated before (Zhao et al., 2018a). These downstream targets of XBP1s transcription factor were not selected randomly; they have certain role in the event of cellular differentiation. High *SEC63* expression had been shown to have involvement in apoptosis and reduced proliferation in hepatic tumors (Casper et al., 2013); ERDJ4 has been shown to have crucial role in B cell differentiation in the event of hematopoiesis (Fritz and Weaver, 2014). Besides, *DNAJC3* is known to get increased during plasma cells differentiation (Shaffer et al., 2004); the expression of *PDIA3* and *DNAJC3* were shown to get up regulated during T-helper cell differentiation and treatment with IRE1α’s endoribonuclease activity specific inhibitor 4µ8c down regulated their expression via inhibiting *XBP1* mRNA splicing (Pramanik et al., 2017). Our results suggested that expression of all these genes increased in kaempferol treated conditions, and get reversed by pretreatment with STF083010 (Fig. 6B). These results confirm that kaempferol activates XBP1s transcriptional activity by activating the endoribonuclease activity of IRE1α and STF083010 blocks the transcriptional activity induced by kaempferol by inhibiting XBP1 mRNA cleavage (Fig. Supplement 2A). Furthermore, we proceeded to check whether APY29, an activator of IRE1α endoribonuclease activity could induce the differentiation of IMR32 cells similar to kaempferol. The western blot analysis of APY29 treated cell lysates showed an increased expression of neuronal differentiation markers similar to that of kaempferol treated conditions (Fig. 6C). The induction of differentiation by APY29 observed in Neuro2a cells was similar to kaempferol treated conditions. There was a marked increase in the β-III tubulin expression and synaptophysin vesicles observed through immunocytochemistry in Neuro2a cells (Fig. Supplement 2B and C). Taken together, these results suggest that kaempferol activates the IRE1α-XBP1 branch pathway to induce the neuroblastoma differentiation. In addition, we analyzed if IRE1α-XBP1 pathway is involved in inducing neuroblastoma differentiation by other agents like CDDO and ATRA (Chaudhari et al., 2017b). However, we found that pre-incubation of cells with STF083010 did not inhibit differentiation induced by CDDO or ATRA (Fig. Supplement 3A and B) suggesting that IRE1α activation is not the only pathway involved in neuroblastoma differentiation. STF083010 (50 µM) did not have cytotoxic effect in IMR32 cells; pretreatment with STF083010 followed by kaempferol treatment also did not modulate cell death (Fig. 6E) suggesting that cell death induction by kaempferol is independent of IRE1α-XBP1 pathway. However, the immunocytochemistry analysis for the expression of β-III tubulin showed reduced differentiation upon STF083010 pre-treatment, in both kaempferol and APY29 treated conditions (Fig. 6D). And also, there was reduced change in morphology and decrease in the number of cells bearing neurites with STF083010 pre-treatment in the presence of kaempferol or APY29 (Fig. 6F and G). Altogether, these results suggest that STF083010 inhibits differentiation but not the cell death induced by kaempferol and APY29. Therefore, the activation of endoribonuclease activity of IRE1α followed by the XBP1 mRNA splicing holds a crucial role in inducing the differentiation of neuroblastoma cells. This could be the reason for the morphological changes and neurite outgrowths observed earlier in other neuroblastoma cell lines upon treatment with flavonoids (Brown et al., 1998a; Sato et al., 1994a).

**Figure 6:**
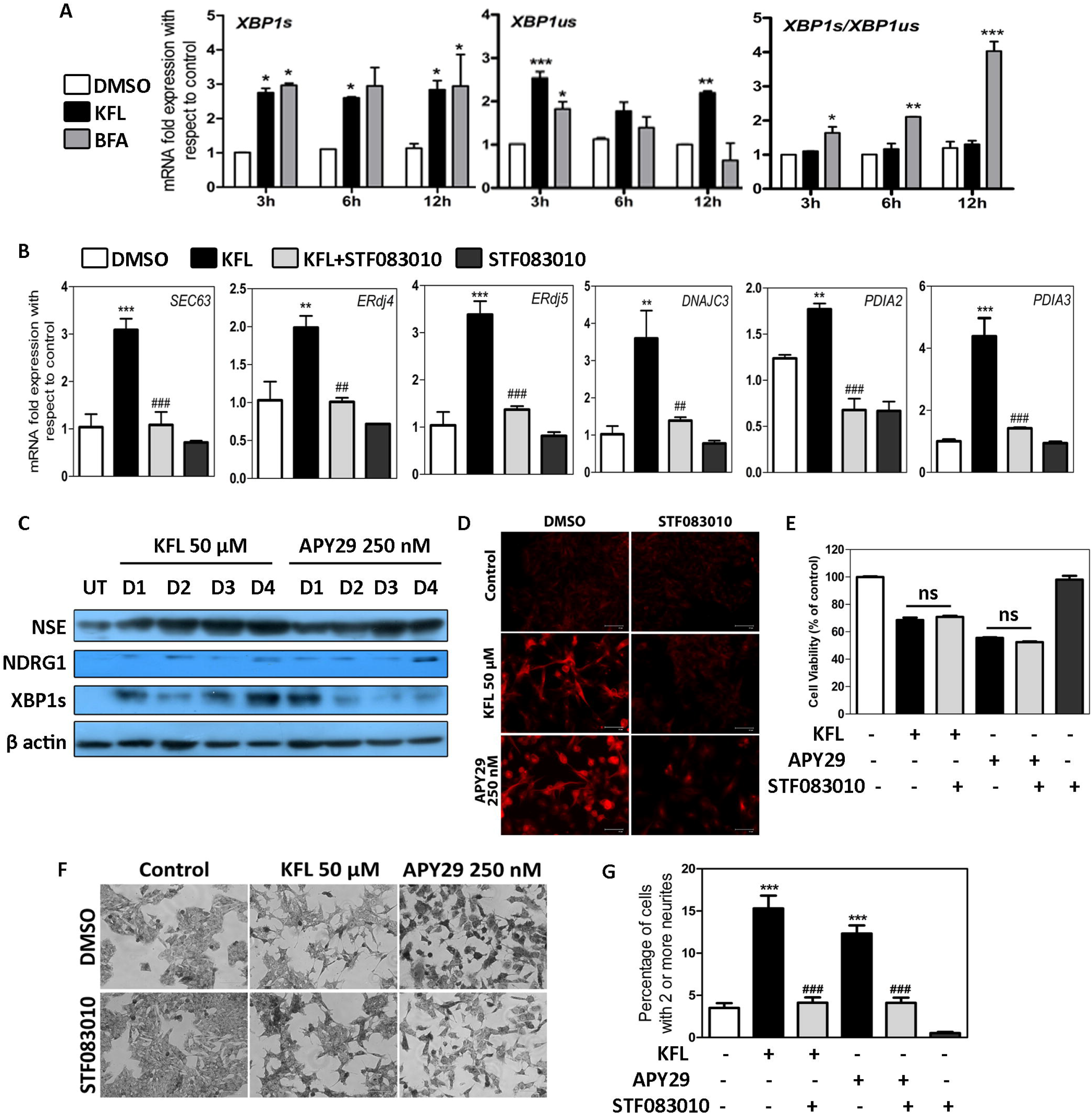
XBP1s expression leads to differentiation of neuroblastoma. **A.** Graphs representing the spliced and unspliced form of xbp1 mRNA present upon kaempferol 50 µM and Brefeldin A (BFA) 1 µg/ml treatment. Ratio represented to point out that kaempferol treated cells maintains high expression levels of xbp1 spliced and unspliced form. The average ±SEM from three independent experiments. (*p < 0.05; **p < 0.01; ***p < 0.001; two-way ANOVA) **B.** Gene expression studies on downstream transcriptional targets of XBP1s in IMR32 cells. Expression levels of mRNA of six downstream targets upon kaempferol (KFL) 50 µM treatment for 12 h was quantified using qRT-PCR. STF083010 was used at 50 µM concentration to reduce the XBP1s activity. The average ±SEM experiments performed in duplicates (n=3). * represents comparison between control and kaempferol treated cells; # represents comparison between kaempferol treated cells with STF083010 treated cells (*p < 0.05; **p < 0.01; ***p < 0.001; one-way ANOVA) **C.** Western blot analysis showing the increase in expression of neuronal marker proteins NSE and NDRG1 with increased XBP1s levels in kaempferol and APY29 treated conditions. **D.** Immunocytochemistry for expression of β III tubulin showing reduced expression upon pretreatment of cells with STF083010 50 µM in the presence of kaempferol and APY29. Representative images of experiments performed more than three times. Scale bar = 67 µm. **E.** Cell death analysis using MTT assay showing no effect of STF083010 on rescuing cell viability upon pretreatment in kaempferol and APY29 treated conditions in IMR32 cells. The average ±SEM from 3 independent experiments performed in triplicates. (*p < 0.05; **p < 0.01; ***p < 0.001; one-way ANOVA) **F.** Phase contrast methylene blue stained images of IMR32 cells showing change in morphology and neurite outgrowth upon treatment with kaempferol (KFL) and APY29 for 96 h. Pretreatment with STF083010 (50 µM) inhibits the process of neuritogenesis. Representative images of experiments performed more than three times. Scale bar = 100 µm. **G.** Graphs representing the number of cells with 2 or more neurite per cell in IMR32 cells after 96 h incubation with kaempferol and APY29. STF083010 pretreatment reduces the number of neurite bearing cells in both kaempferol (KFL) and APY29 treated conditions. The average ±SEM from 10 independent counting is shown.* represents comparison between control and kaempferol/APY29 treated cells; # represents comparison between kaempferol/APY29 treated cells with STF083010 pretreated cells (*p < 0.05; **p < 0.01; ***p < 0.001; one-way ANOVA).

**Figure 7:**
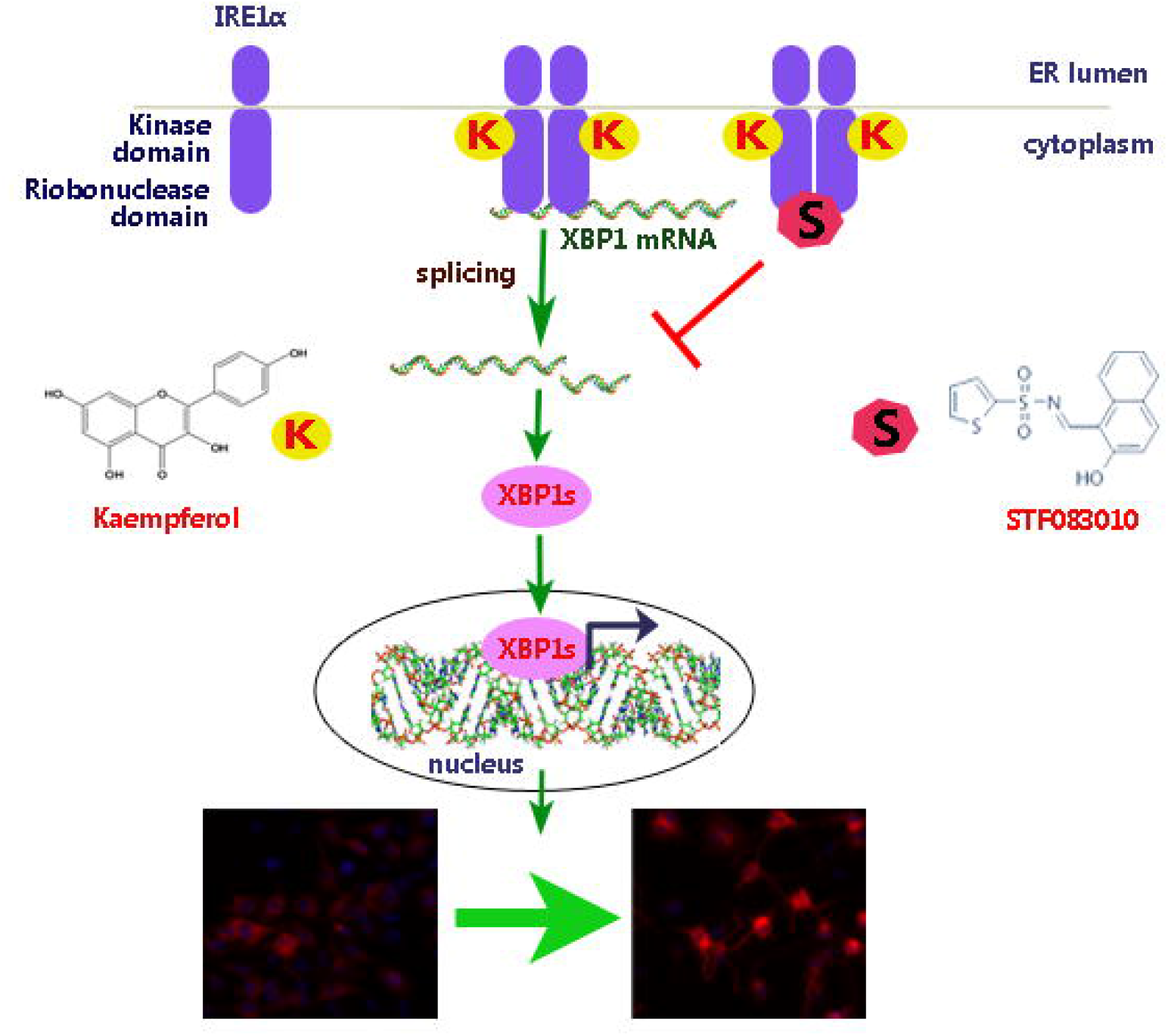
Proposed mechanism of action. Binding of kaempferol (K) to the kinase domain of IRE1α activates its endoribonuclease activity. The transcriptional activity of XBP1 spliced form of protein induces the differentiation of neuroblastoma cells. Pre-treatment with STF083010 (S), an inhibitor for IRE1 ribonuclease activity, inhibits the process of cleavage of XBP1 mRNA thereby inhibits differentiation.

## 4. Discussion

Although several studies have been done in improving the prognosis of high risk neuroblastoma with differentiation therapies, the survival rate of patients remains considerably poor (Berthold et al., 2005; Matthay et al., 2009; Simon et al., 2011). The implementation of chemotherapeutic approaches with synthetic molecules also adds up to the detrimental effects on the patients. Therefore an enhanced understanding of differentiation pathways and novel therapeutic approaches are needed. An approach using dietary compounds or plant metabolites could be a better alternative for reducing the damaging effects of synthetic drugs. Retinoic acid and its derivatives are well explored for their effects in inducing differentiation of neuroblastoma cells in clinical therapies. Other than RA, there are reports on the activity of 17β-estradiol in enhancing the neuritogenesis in neuroblastoma cells, *in vitro* (Mérot et al., 2005). Flavonoids, being known for their anticancer activity, mimic estrogen in physiological systems, while some of them have been reported to induce neuritogenesis in neuroblastoma cells *in vitro* (Brown et al., 1998b; Sato et al., 1994b). Kaempferol had been reported before for its anticancer activity via multiple cellular pathways which includes triggering of ER stress, increasing the expression of DR5 receptor, via estrogen receptor expression and suppressing signal regulated kinases (Guo et al., 2017; Luo et al., 2012; Wang et al., 2013; Zhao et al., 2017).

The role of IRE1α in differentiation of multiple cell types shows its involvement in deciding the fate of cells. *XBP1* mRNA, the major substrate for the IRE1α endoribonuclease activity, had been shown to transcribe genes that are involved in multiple cellular processes, including the genes involved in differentiation of numerous cell types (Cho et al., 2013; Iwakoshi et al., 2003; Reimold et al., 2001b). The role of quercetin, a flavonoid, in activating the endoribonuclease activity of IRE1α gave hint on exploring the anticancer activity of kaempferol in the aspects of IRE1α-XBP1 pathway in the differentiation of neuroblastoma cells (Wiseman et al., 2010). Moreover the inactivation of IRE1α endoribonuclease activity and the hypomorphic expression of XBP1 in intestinal stem cells had been shown to be the reason for intestinal tumorogenesis (Niederreiter et al., 2013).

The difference in the gene expression pattern between kaempferol and ATRA treated conditions shows that they both could activate different pathways for inducing neuroblastoma differentiation. CDDO a triterpenoid was shown to induce differentiation of IMR32 cells via PPAR signalling pathway while ATRA via CREB dependent pathways (Chaudhari et al., 2017a). The difference between the mRNA and protein expression of NSE observed in the ATRA treated condition had been observed in other studies as well. The differentiation of SK-N-DZ cells by overexpressing the Aryl hydrocarbon receptor (AHR) showed no change in the regulation of NSE at transcript levels (Wu et al., 2014). The studies using retinoic acid in IMR32 cell line showed down regulation of NSE mRNA levels at day 5 of differentiation(Maresca et al., 2012). These data suggests that the transcripts of neuronal markers might be regulated via post-transcriptional regulatory mechanisms. Apart from the activation of IRE1α-XPB1 pathway, critical role of activation of other signalling pathways such as cMAP and P38 MAPK pathway (Monaghan et al., 2008), JNK-MAPK pathway and PPAR signalling pathway (Chaudhari et al., 2017a) have been reported to have a critical role in inducing the differentiation of neuroblastoma cells.

In summary, the current study demonstrated the involvement of IRE1α in differentiation of the neuroblastoma cells. The modulation of IRE1α in neuroblastoma cells by kaempferol or by specific modulator entitled for activating the endoribonuclease activity of IRE1α showed enhanced cell death, proliferation repression coupled to differentiation with increase in neuronal marker expressions, an expected outcome for neuroblastoma therapies. The results presented herein demonstrate that kaempferol induces differentiation and cell death of neuroblastoma cells. Our study reporting the differentiation and cell death elicited by kaempferol might suggest new therapeutic approach via modulation of IRE1α for the treatment of neuroblastoma, the infuriating childhood malignancy.

## Acknowledgements

PR greatly acknowledges DST-SERB for the financial support through the Research Grant-SB/EMEQ-223/2014. We thank Ms. Ida Florance PS for her kind help in counting cells bearing neurites.

## Author Contributions

P.R. designed the experiments and A.A. performed all the experiments. Data analysis and manuscript preparation was done by P.R., A.A and PT.

## Conflict of interest

No potential conflicts of interests were disclosed by the authors.

## Supplementary figure legends

**Fig. S1: Kaempferol reduces proliferation of IMR32 cells**

**A.** Trypan blue dye exclusion assay in kaempferol treated cells showing reduced proliferation associated with cell death in 50 and 100 µM concentrations. Data represented as average ±SEM of counting obtained from three independent experiments performed in triplicates (***p < 0.001; two-way ANOVA). Dashed line represents the seeding density of 50×10^3^ cells on day 0.

**B.** Phase contrast images showing the reduction in confluence and change in morphology with kaempferol (KFL) 50 µM with ATRA 25 µM as positive control. Representative image of experiments performed more than three times. Images were acquired at 200x magnification.

**Fig. S2: Modulation of IRE1α endoribonuclease activity by kaempferol**

**A.** Expression of *XBP1s* and *XBP1us* at mRNA level induced by kaempferol (50 µM) in the presence of STF083010 (50 µM) at 12 h. The average ±SEM of three independent experiments. * represents comparison between control and kaempferol treated cells; # represents comparison between kaempferol treated cells with STF083010 treated cells (*p < 0.05; **p < 0.01; ***p < 0.001; one-way ANOVA)

**B & C.** Immunocytochemistry for the expression of β III tubulin and synaptophysin in Neuro2a cells upon treatment with kaempferol (KFL) and APY29 for 96 h. Representative images of experiments performed more than three times. Scale bars = 67 µm.

**Fig. S3: STF083010 inhibition of neuroblastoma differentiation is specific for kaempferol**

**A.** Phase contrast images of IMR32 cells treated with ATRA and CDDO in the presence or absence of STF083010 (50 µM) for a period of 5 days. Representative images of experiment performed for three times. Scale bar = 100µm.

**B.** Graphs representing the number of cells with 2 or more neurite per cell in IMR32 cells after 5 days of incubation with ATRA (25 µM) and CDDO (500nM). STF083010 (50 µM) pretreatment did not reduce the number of neurite bearing cells in both ATRA and CDDO treated conditions. The average ±SEM from five independent counting is shown (non-significant at p < 0.05; one-way ANOVA).

## References

Abdullah, A., Ravanan, P., 2018a. Kaempferol mitigates Endoplasmic Reticulum Stress Induced Cell Death by targeting caspase 3/7. Sci Rep 8, 2189.

Abdullah, A., Ravanan, P., 2018b. The unknown face of IRE1alpha-Beyond ER stress. Eur J Cell Biol 97, 359–368.

Acosta-Alvear, D., Zhou, Y., Blais, A., Tsikitis, M., Lents, N.H., Arias, C., Lennon, C.J., Kluger, Y., Dynlacht, B.D., 2007a. XBP1 controls diverse cell type- and condition-specific transcriptional regulatory networks. Mol Cell 27, 53–66.

Acosta-Alvear, D., Zhou, Y., Blais, A., Tsikitis, M., Lents, N.H., Arias, C., Lennon, C.J., Kluger, Y., Dynlacht, B.D., 2007b. XBP1 controls diverse cell type-and condition-specific transcriptional regulatory networks. Molecular cell 27, 53–66.

Balasubramaniyan, V., Boddeke, E., Bakels, R., Kust, B., Kooistra, S., Veneman, A., Copray, S., 2006. Effects of histone deacetylation inhibition on neuronal differentiation of embryonic mouse neural stem cells. Neuroscience 143, 939–951.

Berthold, F., Boos, J., Burdach, S., Erttmann, R., Henze, G., Hermann, J., Klingebiel, T., Kremens, B., Schilling, F.H., Schrappe, M., 2005. Myeloablative megatherapy with autologous stem-cell rescue versus oral maintenance chemotherapy as consolidation treatment in patients with high-risk neuroblastoma: a randomised controlled trial. The lancet oncology 6, 649–658.

Bork, P., Sander, C., 1993. A hybrid protein kinase-RNase in an interferon-induced pathway? FEBS letters 334, 149–152.

Brodeur, G.M., 2003. Neuroblastoma: biological insights into a clinical enigma. Nat Rev Cancer 3, 203–216.

Brown, A., Jolly, P., Wei, H., 1998a. Genistein modulates neuroblastoma cell proliferation and differentiation through induction of apoptosis and regulation of tyrosine kinase activity and N-myc expression. Carcinogenesis 19, 991–997.

Brown, A., Jolly, P., Wei, H., 1998b. Genistein modulates neuroblastoma cell proliferation and differentiation through induction of apoptosis and regulation of tyrosine kinase activity and N-myc expression. Carcinogenesis 19, 991–997.

Casper, M., Weber, S.N., Kloor, M., Müllenbach, R., Grobholz, R., Lammert, F., Zimmer, V., 2013. Hepatocellular carcinoma as extracolonic manifestation of Lynch syndrome indicates SEC63 as potential target gene in hepatocarcinogenesis. Scandinavian journal of gastroenterology 48, 344–351.

Chaudhari, N., Talwar, P., Lefebvre D’hellencourt, C., Ravanan, P., 2017a. CDDO and ATRA Instigate Differentiation of IMR32 Human Neuroblastoma Cells. Front Mol Neurosci 10, 310.

Chaudhari, N., Talwar, P., Lefebvre D’hellencourt, C., Ravanan, P., 2017b. CDDO and ATRA Instigate Differentiation of IMR32 Human Neuroblastoma Cells. Frontiers in molecular neuroscience 10, 310.

Cho, Y.M., Kim, D.H., Kwak, S.-N., Jeong, S.-W., Kwon, O.-J., 2013. X-box binding protein 1 enhances adipogenic differentiation of 3T3-L1 cells through the downregulation of Wnt10b expression. FEBS letters 587, 1644–1649.

Choi, S.H., Wright, J.B., Gerber, S.A., Cole, M.D., 2010. Myc protein is stabilized by suppression of a novel E3 ligase complex in cancer cells. Genes Dev 24, 1236–1241.

Chow, J.M., Cheng, A.L., Su, I.J., Wang, C.H., 1991. 13-cis-retinoic acid induces cellular differentiation and durable remission in refractory cutaneous ki-1 lymphoma. Cancer 67, 2490–2494.

Concha, N.O., Smallwood, A., Bonnette, W., Totoritis, R., Zhang, G., Federowicz, K., Yang, J., Qi, H., Chen, S., Campobasso, N., 2015. Long-range inhibitor-induced conformational regulation of human IRE1α endoribonuclease activity. Molecular pharmacology 88, 1011–1023.

Cui, Y., Morgenstern, H., Greenland, S., Tashkin, D.P., Mao, J.T., Cai, L., Cozen, W., Mack, T.M., Lu, Q.Y., Zhang, Z.F., 2008. Dietary flavonoid intake and lung cancer--a population-based case-control study. Cancer 112, 2241–2248.

Duckett, J.W., Koop, C.E., 1977. Neuroblastoma. Urol Clin North Am 4, 285–295.

Fritz, J.M., Weaver, T.E., 2014. The BiP cochaperone ERdj4 is required for B cell development and function. PloS one 9, e107473.

Gates, M.A., Vitonis, A.F., Tworoger, S.S., Rosner, B., Titus-Ernstoff, L., Hankinson, S.E., Cramer, D.W., 2009. Flavonoid intake and ovarian cancer risk in a population-based case-control study. Int J Cancer 124, 1918–1925.

Guglielmi, L., Cinnella, C., Nardella, M., Maresca, G., Valentini, A., Mercanti, D., Felsani, A., D’Agnano, I., 2014. MYCN gene expression is required for the onset of the differentiation programme in neuroblastoma cells. Cell Death Dis 5, e1081.

Guo, H., Lin, W., Zhang, X., Zhang, X., Hu, Z., Li, L., Duan, Z., Zhang, J., Ren, F., 2017. Kaempferol induces hepatocellular carcinoma cell death via endoplasmic reticulum stress-CHOP-autophagy signaling pathway. Oncotarget 8, 82207.

Guo, H., Ren, F., Zhang, L., Zhang, X., Yang, R., Xie, B., Li, Z., Hu, Z., Duan, Z., Zhang, J., 2016. Kaempferol induces apoptosis in HepG2 cells via activation of the endoplasmic reticulum stress pathway. Molecular medicine reports 13, 2791–2800.

Halder, D., Kim, G.H., Shin, I., 2015. Synthetic small molecules that induce neuronal differentiation in neuroblastoma and fibroblast cells. Mol Biosyst 11, 2727–2737.

Han, D., Lerner, A.G., Walle, L.V., Upton, J.-P., Xu, W., Hagen, A., Backes, B.J., Oakes, S.A., Papa, F.R., 2009. IRE1α kinase activation modes control alternate endoribonuclease outputs to determine divergent cell fates. Cell 138, 562–575.

Hoehner, J.C., Gestblom, C., Hedborg, F., Sandstedt, B., Olsen, L., Pahlman, S., 1996. A developmental model of neuroblastoma: differentiating stroma-poor tumors’ progress along an extra-adrenal chromaffin lineage. Lab Invest 75, 659–675.

Huang, W.W., Chiu, Y.J., Fan, M.J., Lu, H.F., Yeh, H.F., Li, K.H., Chen, P.Y., Chung, J.G., Yang, J.S., 2010. Kaempferol induced apoptosis via endoplasmic reticulum stress and mitochondria-dependent pathway in human osteosarcoma U-2 OS cells. Molecular nutrition & food research 54, 1585–1595.

Iwakoshi, N.N., Lee, A.H., Vallabhajosyula, P., Otipoby, K.L., Rajewsky, K., Glimcher, L.H., 2003. Plasma cell differentiation and the unfolded protein response intersect at the transcription factor XBP-1. Nat Immunol 4, 321–329.

Iwawaki, T., Akai, R., Yamanaka, S., Kohno, K., 2009. Function of IRE1 alpha in the placenta is essential for placental development and embryonic viability. Proc Natl Acad Sci U S A 106, 16657–16662.

Jones-Villeneuve, E., McBURNEY, M.W., Rogers, K.A., Kalnins, V.I., 1982. Retinoic acid induces embryonal carcinoma cells to differentiate into neurons and glial cells. The Journal of cell biology 94, 253–262.

Kang, J.H., Rychahou, P.G., Ishola, T.A., Qiao, J., Evers, B.M., Chung, D.H., 2006. MYCN silencing induces differentiation and apoptosis in human neuroblastoma cells. Biochem Biophys Res Commun 351, 192–197.

Kim, S.-H., Choi, K.-C., 2013. Anti-cancer effect and underlying mechanism (s) of kaempferol, a phytoestrogen, on the regulation of apoptosis in diverse cancer cell models. Toxicological research 29, 229.

Kollareddy, M., Sherrard, A., Park, J.H., Szemes, M., Gallacher, K., Melegh, Z., Oltean, S., Michaelis, M., Cinatl, J., Jr., Kaidi, A., Malik, K., 2017. The small molecule inhibitor YK-4–279 disrupts mitotic progression of neuroblastoma cells, overcomes drug resistance and synergizes with inhibitors of mitosis. Cancer Lett 403, 74–85.

Korennykh, A.V., Egea, P.F., Korostelev, A.A., Finer-Moore, J., Zhang, C., Shokat, K.M., Stroud, R.M., Walter, P., 2009. The unfolded protein response signals through high-order assembly of Ire1. Nature 457, 687.

Kuiper, G.G., Lemmen, J.G., Carlsson, B., Corton, J.C., Safe, S.H., Van Der Saag, P.T., Van Der Burg, B., Gustafsson, J.-A.k., 1998. Interaction of estrogenic chemicals and phytoestrogens with estrogen receptor β. Endocrinology 139, 4252–4263.

Lange, C., Mix, E., Frahm, J., Glass, Ä., Müller, J., Schmitt, O., Schmöle, A.-C., Klemm, K., Ortinau, S., Hübner, R., 2011. Small molecule GSK-3 inhibitors increase neurogenesis of human neural progenitor cells. Neuroscience letters 488, 36–40.

Lee, K.P., Dey, M., Neculai, D., Cao, C., Dever, T.E., Sicheri, F., 2008. Structure of the dual enzyme Ire1 reveals the basis for catalysis and regulation in nonconventional RNA splicing. Cell 132, 89–100.

Lu, J., Guan, S., Zhao, Y., Yu, Y., Woodfield, S.E., Zhang, H., Yang, K.L., Bieerkehazhi, S., Qi, L., Li, X., Gu, J., Xu, X., Jin, J., Muscal, J.A., Yang, T., Xu, G.T., Yang, J., 2017. The second-generation ALK inhibitor alectinib effectively induces apoptosis in human neuroblastoma cells and inhibits tumor growth in a TH-MYCN transgenic neuroblastoma mouse model. Cancer Lett 400, 61–68.

Luo, H., Rankin, G.O., Juliano, N., Jiang, B.-H., Chen, Y.C., 2012. Kaempferol inhibits VEGF expression and in vitro angiogenesis through a novel ERK-NFκB-cMyc-p21 pathway. Food chemistry 130, 321–328.

Ma, Y., Shimizu, Y., Mann, M.J., Jin, Y., Hendershot, L.M., 2010. Plasma cell differentiation initiates a limited ER stress response by specifically suppressing the PERK-dependent branch of the unfolded protein response. Cell Stress and Chaperones 15, 281–293.

Maresca, G., Natoli, M., Nardella, M., Arisi, I., Trisciuoglio, D., Desideri, M., Brandi, R., D’Aguanno, S., Nicotra, M.R., D’Onofrio, M., Urbani, A., Natali, P.G., Del Bufalo, D., Felsani, A., D’Agnano, I., 2012. LMNA knock-down affects differentiation and progression of human neuroblastoma cells. PLoS One 7, e45513.

Matsuzaki, S., Hiratsuka, T., Taniguchi, M., Shingaki, K., Kubo, T., Kiya, K., Fujiwara, T., Kanazawa, S., Kanematsu, R., Maeda, T., 2015. Physiological ER stress mediates the differentiation of fibroblasts. PloS one 10, e0123578.

Matthay, K.K., Reynolds, C.P., Seeger, R.C., Shimada, H., Adkins, E.S., Haas-Kogan, D., Gerbing, R.B., London, W.B., Villablanca, J.G., 2009. Long-term results for children with high-risk neuroblastoma treated on a randomized trial of myeloablative therapy followed by 13-cis-retinoic acid: a children’s oncology group study. Journal of clinical oncology 27, 1007.

Matthay, K.K., Villablanca, J.G., Seeger, R.C., Stram, D.O., Harris, R.E., Ramsay, N.K., Swift, P., Shimada, H., Black, C.T., Brodeur, G.M., Gerbing, R.B., Reynolds, C.P., 1999. Treatment of high-risk neuroblastoma with intensive chemotherapy, radiotherapy, autologous bone marrow transplantation, and 13-cis-retinoic acid. Children’s Cancer Group. N Engl J Med 341, 1165–1173.

McCann, S.E., Ambrosone, C.B., Moysich, K.B., Brasure, J., Marshall, J.R., Freudenheim, J.L., Wilkinson, G.S., Graham, S., 2005. Intakes of selected nutrients, foods, and phytochemicals and prostate cancer risk in western New York. Nutr Cancer 53, 33–41.

Mérot, Y., Ferrière, F., Debroas, E., Flouriot, G., Duval, D., Saligaut, C., 2005. Estrogen receptor alpha mediates neuronal differentiation and neuroprotection in PC12 cells: critical role of the A/B domain of the receptor. Journal of molecular endocrinology 35, 257–267.

Monaghan, T.K., Mackenzie, C.J., Plevin, R., Lutz, E.M., 2008. PACAP-38 induces neuronal differentiation of human SH-SY5Y neuroblastoma cells via cAMP-mediated activation of ERK and p38 MAP kinases. J Neurochem 104, 74–88.

Niederreiter, L., Fritz, T.M., Adolph, T.E., Krismer, A.M., Offner, F.A., Tschurtschenthaler, M., Flak, M.B., Hosomi, S., Tomczak, M.F., Kaneider, N.C., Sarcevic, E., Kempster, S.L., Raine, T., Esser, D., Rosenstiel, P., Kohno, K., Iwawaki, T., Tilg, H., Blumberg, R.S., Kaser, A., 2013. ER stress transcription factor Xbp1 suppresses intestinal tumorigenesis and directs intestinal stem cells. J Exp Med 210, 2041–2056.

Niwa, M., Patil, C.K., DeRisi, J., Walter, P., 2005. Genome-scale approaches for discovering novel nonconventional splicing substrates of the Ire1 nuclease. Genome Biol 6, R3.

Papandreou, I., Denko, N.C., Olson, M., Van Melckebeke, H., Lust, S., Tam, A., Solow-Cordero, D.E., Bouley, D.M., Offner, F., Niwa, M., Koong, A.C., 2011. Identification of an Ire1alpha endonuclease specific inhibitor with cytotoxic activity against human multiple myeloma. Blood 117, 1311–1314.

Pemrick, S.M., Lucas, D., Grippo, J., 1994. The retinoid receptors. Leukemia 8, S1–10.

Pence, J.C., Shorter, N.A., 1990. In vitro differentiation of human neuroblastoma cells caused by vasoactive intestinal peptide. Cancer research 50, 5177–5183.

Pramanik, J., Chen, X., Kar, G., Gomes, T., Henriksson, J., Miao, Z., Natarajan, K., McKenzie, A.N., Mahata, B., Teichmann, S.A., 2017. The IRE1a-XBP1 pathway promotes T helper cell differentiation by resolving secretory stress and accelerating proliferation. bioRxiv, 235010.

Ravanan, P., Sano, R., Talwar, P., Ogasawara, S., Matsuzawa, S.-i., Cuddy, M., Singh, S.K., Rao, G.S., Kondaiah, P., Reed, J.C., 2011. Synthetic triterpenoid cyano enone of methyl boswellate activates intrinsic, extrinsic, and endoplasmic reticulum stress cell death pathways in tumor cell lines. Molecular cancer therapeutics 10, 1635–1643.

Reimold, A.M., Iwakoshi, N.N., Manis, J., Vallabhajosyula, P., Szomolanyi-Tsuda, E., Gravallese, E.M., Friend, D., Grusby, M.J., Alt, F., Glimcher, L.H., 2001a. Plasma cell differentiation requires the transcription factor XBP-1. Nature 412, 300–307.

Reimold, A.M., Iwakoshi, N.N., Manis, J., Vallabhajosyula, P., Szomolanyi-Tsuda, E., Gravallese, E.M., Friend, D., Grusby, M.J., Alt, F., Glimcher, L.H., 2001b. Plasma cell differentiation requires the transcription factor XBP-1. Nature 412, 300.

Rettig, I., Koeneke, E., Trippel, F., Mueller, W.C., Burhenne, J., Kopp-Schneider, A., Fabian, J., Schober, A., Fernekorn, U., von Deimling, A., Deubzer, H.E., Milde, T., Witt, O., Oehme, I., 2015. Selective inhibition of HDAC8 decreases neuroblastoma growth in vitro and in vivo and enhances retinoic acid-mediated differentiation. Cell Death Dis 6, e1657.

Sato, F., Matsukawa, Y., Matsumoto, K., Nishino, H., Sakai, T., 1994a. Apigenin induces morphological differentiation and G2-M arrest in rat neuronal cells. Biochemical and biophysical research communications 204, 578–584.

Sato, F., Matsukawa, Y., Matsumoto, K., Nishino, H., Sakai, T., 1994b. Apigenin induces morphological differentiation and G2-M arrest in rat neuronal cells. Biochem Biophys Res Commun 204, 578–584.

Sha, H., He, Y., Chen, H., Wang, C., Zenno, A., Shi, H., Yang, X., Zhang, X., Qi, L., 2009a. The IRE1alpha-XBP1 pathway of the unfolded protein response is required for adipogenesis. Cell Metab 9, 556–564.

Sha, H., He, Y., Chen, H., Wang, C., Zenno, A., Shi, H., Yang, X., Zhang, X., Qi, L., 2009b. The IRE1α-XBP1 pathway of the unfolded protein response is required for adipogenesis. Cell metabolism 9, 556–564.

Shaffer, A., Shapiro-Shelef, M., Iwakoshi, N.N., Lee, A.-H., Qian, S.-B., Zhao, H., Yu, X., Yang, L., Tan, B.K., Rosenwald, A., 2004. XBP1, downstream of Blimp-1, expands the secretory apparatus and other organelles, and increases protein synthesis in plasma cell differentiation. Immunity 21, 81–93.

Simon, T., Berthold, F., Borkhardt, A., Kremens, B., De Carolis, B., Hero, B., 2011. Treatment and outcomes of patients with relapsed, high-risk neuroblastoma: Results of German trials. Pediatric blood & cancer 56, 578–583.

Takahashi, J., Palmer, T.D., Gage, F.H., 1999. Retinoic acid and neurotrophins collaborate to regulate neurogenesis in adult-derived neural stem cell cultures. Developmental Neurobiology 38, 65–81.

Takahashi, N., Koyama, S., Hasegawa, S., Yamasaki, M., Imai, M., 2017. Anticancer efficacy of p-dodecylaminophenol against high-risk and refractory neuroblastoma cells in vitro and in vivo. Bioorg Med Chem Lett 27, 4664–4672.

Todd, D.J., McHeyzer-Williams, L.J., Kowal, C., Lee, A.-H., Volpe, B.T., Diamond, B., McHeyzer-Williams, M.G., Glimcher, L.H., 2009. XBP1 governs late events in plasma cell differentiation and is not required for antigen-specific memory B cell development. Journal of Experimental Medicine 206, 2151–2159.

Tohmonda, T., Miyauchi, Y., Ghosh, R., Yoda, M., Uchikawa, S., Takito, J., Morioka, H., Nakamura, M., Iwawaki, T., Chiba, K., Toyama, Y., Urano, F., Horiuchi, K., 2011. The IRE1alpha-XBP1 pathway is essential for osteoblast differentiation through promoting transcription of Osterix. EMBO Rep 12, 451–457.

Tohmonda, T., Yoda, M., Mizuochi, H., Morioka, H., Matsumoto, M., Urano, F., Toyama, Y., Horiuchi, K., 2013. The IRE1α-XBP1 pathway positively regulates parathyroid hormone (PTH)/PTH-related peptide receptor expression and is involved in pth-induced osteoclastogenesis. Journal of Biological Chemistry 288, 1691–1695.

Tsuru, A., Imai, Y., Saito, M., Kohno, K., 2016. Novel mechanism of enhancing IRE1α-XBP1 signalling via the PERK-ATF4 pathway. Scientific reports 6, 24217.

Urano, F., Wang, X., Bertolotti, A., Zhang, Y., Chung, P., Harding, H.P., Ron, D., 2000. Coupling of stress in the ER to activation of JNK protein kinases by transmembrane protein kinase IRE1. Science 287, 664–666.

Wang, H., Gao, M., Wang, J., 2013. Kaempferol inhibits cancer cell growth by antagonizing estrogen-related receptor α and γ activities. Cell biology international 37, 1190–1196.

Wang, J., Fang, F., Huang, Z., Wang, Y., Wong, C., 2009. Kaempferol is an estrogen-related receptor alpha and gamma inverse agonist. FEBS Lett 583, 643–647.

Wang, J., Fang, X., Ge, L., Cao, F., Zhao, L., Wang, Z., Xiao, W., 2018. Antitumor, antioxidant and anti-inflammatory activities of kaempferol and its corresponding glycosides and the enzymatic preparation of kaempferol. PLoS One 13, e0197563.

Wang, L., Perera, B.G.K., Hari, S.B., Bhhatarai, B., Backes, B.J., Seeliger, M.A., Schürer, S.C., Oakes, S.A., Papa, F.R., Maly, D.J., 2012. Divergent allosteric control of the IRE1α endoribonuclease using kinase inhibitors. Nature chemical biology 8, 982–989.

Watabe, H., Soma, Y., Ito, M., Kawa, Y., Mizoguchi, M., 2002. All-trans Retinoic Acid Induces Differentiation and Apoptosis of Murine Melanocyte Precursors with Induction of the Microphthalmia-Associated Transcription Factor. Journal of Investigative Dermatology 118, 35–42.

Wichterle, H., Lieberam, I., Porter, J.A., Jessell, T.M., 2002. Directed differentiation of embryonic stem cells into motor neurons. Cell 110, 385–397.

Wielenga, M.C., Colak, S., Heijmans, J., de Jeude, J.F.v.L., Rodermond, H.M., Paton, J.C., Paton, A.W., Vermeulen, L., Medema, J.P., van den Brink, G.R., 2015. ER-stress-induced differentiation sensitizes colon cancer stem cells to chemotherapy. Cell reports 13, 489–494.

Wiseman, R.L., Zhang, Y., Lee, K.P., Harding, H.P., Haynes, C.M., Price, J., Sicheri, F., Ron, D., 2010. Flavonol activation defines an unanticipated ligand-binding site in the kinase-RNase domain of IRE1. Molecular cell 38, 291–304.

Wu, P.Y., Liao, Y.F., Juan, H.F., Huang, H.C., Wang, B.J., Lu, Y.L., Yu, I.S., Shih, Y.Y., Jeng, Y.M., Hsu, W.M., Lee, H., 2014. Aryl hydrocarbon receptor downregulates MYCN expression and promotes cell differentiation of neuroblastoma. PLoS One 9, e88795.

Zhang, W., Zeng, Y.-S., Zhang, X.-B., Wang, J.-M., Chen, S.-J., 2006. Combination of adenoviral vector-mediated neurotrophin-3 gene transfer and retinoic acid promotes adult bone marrow cells to differentiate into neuronal phenotypes. Neuroscience letters 408, 98–103.

Zhao, N., Cao, J., Xu, L., Tang, Q., Dobrolecki, L.E., Lv, X., Talukdar, M., Lu, Y., Wang, X., Hu, D.Z., 2018a. Pharmacological targeting of MYC-regulated IRE1/XBP1 pathway suppresses MYC-driven breast cancer. The Journal of clinical investigation 128.

Zhao, P., Aguilar, A.E., Lee, J.Y., Paul, L.A., Suh, J.H., Puri, L., Zhang, M., Beckstead, J., Witkowski, A., Ryan, R.O., Saba, J.D., 2018b. Sphingadienes show therapeutic efficacy in neuroblastoma in vitro and in vivo by targeting the AKT signaling pathway. Invest New Drugs.

Zhao, Y., Tian, B., Wang, Y., Ding, H., 2017. Kaempferol Sensitizes Human Ovarian Cancer Cells-OVCAR-3 and SKOV-3 to Tumor Necrosis Factor-Related Apoptosis-Inducing Ligand (TRAIL)-Induced Apoptosis via JNK/ERK-CHOP Pathway and Up-Regulation of Death Receptors 4 and 5. Medical science monitor: international medical journal of experimental and clinical research 23, 5096.

